# Video-driven simulation of lower limb mechanical loading during aquatic exercises

**DOI:** 10.1101/2022.11.23.517406

**Authors:** Jessy Lauer

## Abstract

Understanding the mechanical demands of an exercise on the musculoskeletal system is crucial to prescribe effective training or therapeutic interventions. Yet, that knowledge is currently limited in water, mostly because of the difficulty in evaluating external resistance. Here I reconcile recent advances in 3D markerless pose and mesh estimation, biomechanical simulations, and hydrodynamic modeling, to predict lower limb mechanical loading during aquatic exercises. Simulations are driven exclusively from a single video. In silico hip and knee joint forces agreed well with in vivo instrumented implant recordings downloaded from the OrthoLoad database, both in magnitude and direction. New insights into individual muscle contributions to joint loading were gained. This noninvasive method has the potential to standardize the reporting of exercise intensity, inform the design of rehabilitation protocols and improve their reproducibility.

## Introduction

Aquatic exercise is a widespread and versatile modality of rehabilitation and training. Thanks to its hydrodynamic properties, water confers unique mechanical benefits (e.g., (1)). Notably, buoyancy offloads weight bearing and promotes greater range of motion; viscosity makes water highly dampening, protecting against injury; and pressure provides a formidable source of resistance that is easily modulated by manipulating movement speed or wetted area. Taken together, these make water-based exercise a nice complement to dry-land training in athletes (2), and a therapy effective in improving physical function in healthy older adults (3) and populations with a variety of chronic conditions (such as osteoarthritis (4) or stroke (5)).

Exercise in water may nonetheless be inadequately prescribed. Systematic reviews point out high methodological heterogeneity among studies, and touch on the limited knowledge of an exercise intensity and its effect on muscle function (6–8). Unlike on land, external forces in water are difficult to quantify: they require, among others, the determination of the pressure field over the moving segments, as well as the energy lost into displacing the water around them. Owing to the lack of measurement tools, this impedes not only the evaluation of an exercise’s mechanical demands but also its prescription. For example, aquatic exercise is believed to be often recommended at too low a dose (e.g. (6)), creating stimuli that are likely insufficient to elicit optimal functional gains. To further complicate matters, exercise intensity is underreported and evidence-based guidelines are scarce, hindering the design and reproducibility of exercise protocols (3, 8).

In order to conceive evidence-based, safe, and graded aquatic exercise programs, quantitative estimates of mechanical loading are vital (e.g., (9)). Several studies have used surface electromyography toward this goal, though with modest success (10)). Although valuable in inferring the timing of excitation and (indirectly) exercise resistance (e.g., (11)), myoelectric activity patterns are not suitable to draw conclusions about muscle force production (12). Another, biomechanically informed strategy is to mathematically describe anatomical structures and hydrodynamic resistances and solve for internal load. This had been elegantly done in early works evaluating joint forces on a simplified knee–shank system during underwater knee extensions (13) and during walking in shallow water (14). A more sophisticated approach exists, substituting simple hydrodynamic models with industry-grade numerical fluid flow simulations, in combination with a multibody dynamics formulation that is robust to noise and modeling errors (15, 16). Resolution of fine flow features however comes at a prohibitive computational cost, which is impractical for routine clinical use. Furthermore, despite the unprecedented mechanistic insights gained into the mechanical demands of aquatic exercises, none of these studies truly probed muscle or joint loads. This is because inverse dynamics (the computational framework they used) does not incorporate anatomically defined muscles by design, so their contribution to joint loading is ignored (17); joint forces computed in this way are in effect only partial loads that greatly underestimate structural load (17, 18).

The most accurate and reliable load measurements are arguably recorded in vivo from instrumented implants. For example, lower limb joint loading during non-weight bearing activities was found to be ≈2.7× lower in water than on land; as speed increased (or with the use of a resistive device), changes in peak forces were nonlinear and less pronounced at the knee than at the hip, underlining the complex interplay of hydrodynamics, exercise type, and muscle function (19). A noninvasive solution is musculoskeletal modeling—a computational framework to elucidate the causal relationships between neuromuscular control, body movement, and internal forces (e.g., (20)). Still, it is typically confined to research environments with expensive, permanent motion capture systems, impeding its use in the field (let alone in a swimming pool, where the necessary use of reflective markers substantially increases drag (21)). Recently though, video-based techniques powered by deep learning have offered a remarkably potent alternative to reconstructing 2D and 3D kinematics under more ecological conditions (22). Building on these advances, I describe a markerless method interfacing gold-standard biomechanical simulations with computer vision for the prediction of joint mechanical loading from a monocular video. I validate its use in aquatic exercises via direct comparison with in vivo load recordings across various conditions, and demonstrate its utility towards better understanding muscle function in water and planning exercise programs.

## Methods

The processing pipeline consists of four consecutive steps. As illustrated in Fig 1, a monocular video is first fed into a deep neural network for the extraction of 2D keypoint coordinates. Second, these are lifted to 3D, and mesh vertices are simultaneously estimated. Third, hydrodynamic forces are computed over the discretized body surface using blade-element theory. Lastly, 3D kinematics and external forces are input into scaled musculoskeletal models for the prediction of muscle and joint reaction forces.

**Fig. 1.**
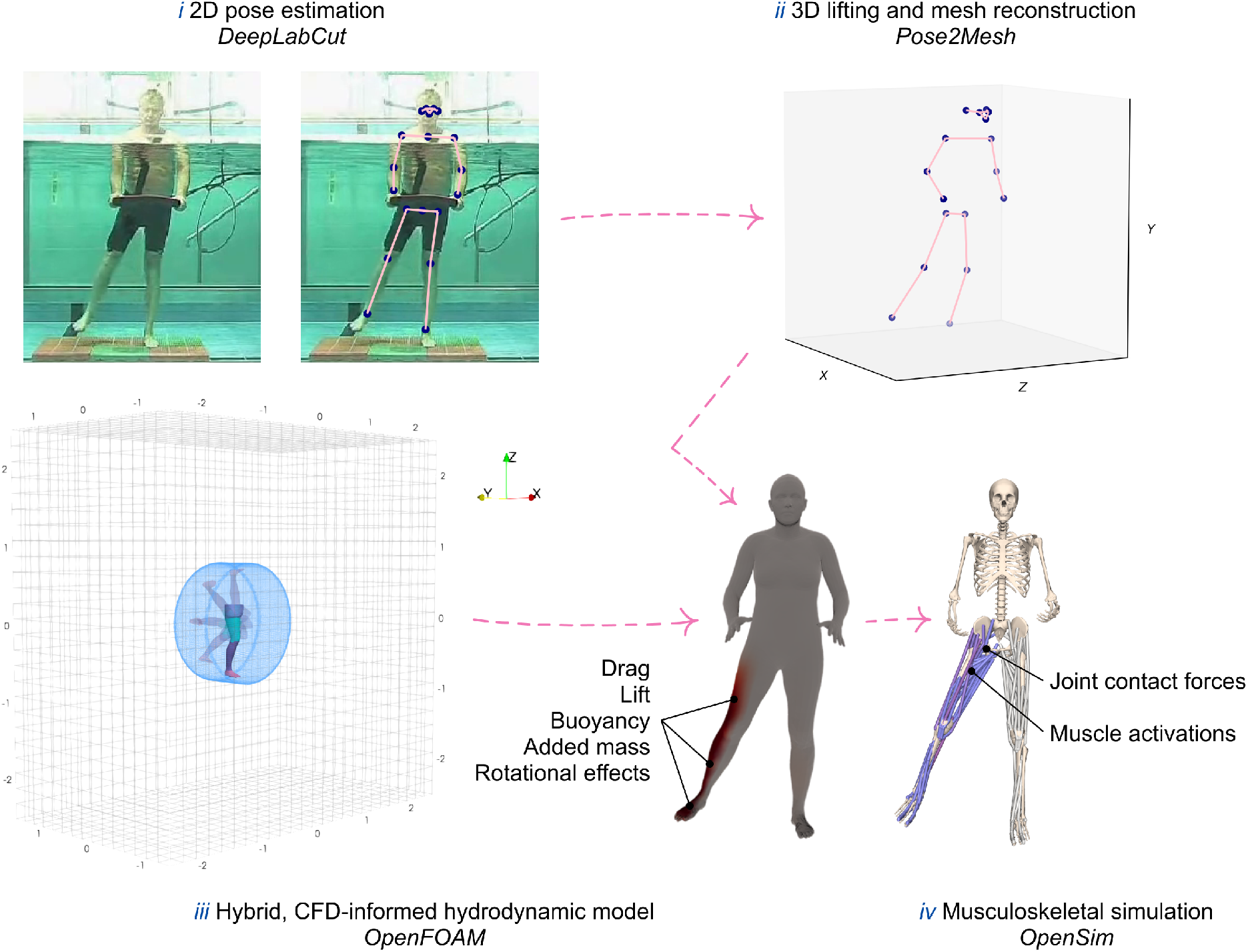
Data processing workflow. (i) Monocular videos from the OrthoLoad database are analyzed using DeepLabCut to predict the 2D coordinates of keypoints with deep learning. (ii) Predicted poses are then fed into Pose2Mesh to lift keypoints to 3D and estimate body meshes. (iii) Drag, lift, and other transient hydrodynamical forces are calculated using a blade element model informed by OpenFOAM’s high-fidelity numerical fluid flow simulations, and integrated over the triangular mesh surface previously obtained. (iv) Last, kinematics, external forces, and scaled full-body musculoskeletal models are input into OpenSim to compute joint contact forces and muscle activations. Note that outputs of musculoskeletal models are not restricted to these two quantities; they can, for example, also include ligament/tendon strains or metabolics.

### Data curation

Video files and in vivo (ground truth) hip and knee joint loads were downloaded from OrthoLoad—a free public database of instrumented implant measurements during routine and sports activities (23). The following search criteria were used: Implant: Hip Joint III or Knee Joint; For aquatic exercises (Parameter: accordingly; Patient: all); Activity: Sports, Aqua Gym, and specifically non-weight bearing dynamic activities; i.e., hip abduction/adduction, hip flexion/extension, knee flexion/extension, and (recumbent) cycling aided with a pool noodle in the back. Regions of 410 × 330 pixels were cropped from the bottom right of the videos so that participants only were visible. Experiments are described in detail in (19). In short, exercises were videotaped at 50 Hz from a lateral window in the pool, and performed in chest-deep water at 35 (slow) and 70 (fast) bpm with the help of acoustic feedback, for four repetitions on average (range: 2–7). Additionally, some were performed with a small resistive device (Aquafins, TheraBand, USA) attached to the ipsilateral ankle. Subjects’ data and the number of videos in which they appear are summarized in Table 1.

**Table 1.** Subjects’ data. Subjects’ codes indicate the type of implant (H: hip; K: knee) and the side (R: right; L: left). m: male; f: female.

| subject | sex | height | mass | age | # exercises |
| --- | --- | --- | --- | --- | --- |
| H2R | m | 1.72 | 84 | 65 | 7 |
| H6R | m | 1.76 | 85 | 71 | 6 |
| H7R | m | 1.79 | 91 | 55 | 7 |
| H8L | m | 1.78 | 94 | 58 | 1 |
| K2L | m | 1.71 | 90 | 79 | 3 |
| K5R | m | 1.75 | 90 | 66 | 5 |
| K7L | f | 1.66 | 63 | 80 | 1 |
| K8L | m | 1.74 | 76 | 76 | 5 |

### 3D pose and mesh estimation

Video-based 2D pose estimation^1^ was done with DeepLabCut v2.2.0.2—a Python toolbox powered by deep learning algorithms available at https://github.com/DeepLabCut/DeepLabCut (25, 26). We used a ResNet-101 network pretrained on the MPII Human Pose dataset (27). To make it generalize well to videos of aquatic exercises, we fine-tuned it using 240 additional frames. These were automatically extracted from 24 videos covering all four activities, and using *k*-means clustering to encourage diversity in their visual content (28). Video frames were manually annotated using DeepLabCut’s GUI, labeling 14 keypoints (left and right ankles, knees, hips, shoulders, elbows, wrists, and chin, and forehead). To assess the robustness of the model to unseen data, annotations from patients H6R and K8L, as well as all fast speed exercises, were left out for testing; i.e., of the 240 images, 110 only were used for training. The network was retrained with a batch size of 16 and data augmentation (scale jitter: 0.5–1.25; rotation: ±25 deg; coarse dropout: *p* = 2%, size = 30%; motion blur: *k* = 7; elastic transformation: 5*σ*) for 50k iterations (after what the loss had plateaued). Network performance was evaluated by calculating the root mean square error between human annotations and DeepLabCut’s predictions, and visually inspecting labeled images.

Three-dimensional poses and meshes were recovered from the 2D poses using Pose2Mesh (29). Pose2Mesh consists of two networks built in a cascaded architecture: PoseNet, which lifts 2D poses to 3D; and MeshNet, which takes both 2D and 3D poses as input and estimates 3D mesh geometry and topology in a model-free manner. As Pose2Mesh relies exclusively on the information contained in the body posture, it does not overfit to image appearance, and thus generalize well to in-the-wild data. To produce smoother mesh sequences, jitter in the input 2D poses was filtered with a 1 C filter (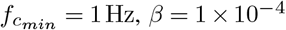 (30)). Unlike other filtering methods, the 1 C filter is adaptive; i.e., it automatically tunes its cutoff frequency according to movement speed, reducing jitter at the expense of lag at low speeds, and vice versa.

#### Hybrid, CFD-informed hydrodynamic modeling

Hydrodynamic forces are computed using blade-element theory (often called strip theory in marine engineering) and quasisteady aerodynamic models. These models, abundantly described in studies of flight analysis, are readily applicable to aquatic locomotion given the similarity of force generation mechanisms in both media (31). Moreover, and in contrast to costly numerical simulations that accurately model flow dynamics, quasi-steady modeling is better suited for the rapid approximation of hydrodynamic forces and torques. The method consists in discretizing the body surface into infinitesimal elements onto which local hydrodynamic forces are evaluated and summed up to determine the net force acting over the entire body. Thus, the 3D triangular meshes predicted by Pose2Mesh can be readily used as input.

#### Calculation of quasi-steady forces

The instantaneous hydrodynamic force acting on the moving lower limb ***F***_*hydro*_ is decomposed into three additive force components: drag ***F***_*d*_ and lift ***F***_*l*_, respectively opposite and perpendicular to the leg’s instantaneous velocity; and added mass ***F***_*am*_, the reaction force caused by the volume of fluid being accelerated together with the leg as its velocity changes:

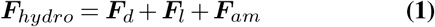

Friction was neglected as it was found to represent < 1.5% of the total force in preliminary numerical simulations. Buoyancy, omitted from the above equation as it is not considered a quasi-steady force, was customarily calculated as: ***F***_*b*_ = *ρV g*, where *g* is the gravitational acceleration, *ρ* is the water density, and *V* is the segment volume determined from the body meshes. Quasi-steady forces are calculated as follows:

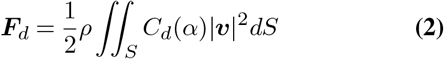

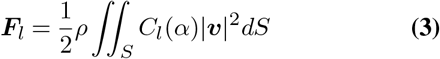

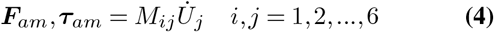

where *C*_*d*_ and *C*_*l*_ are the drag and lift coefficients that are functions of the angle of attack *α* (how they were determined is explained in the next section), ||***v***|| is the magnitude of the face centroid velocity, *dS* is the face area, *M* is the added mass matrix, ***τ***_*am*_ is the torque due to added mass caused by angular accelerations, and 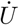 is a 6D vector concatenating linear and angular body accelerations.

For high-Reynolds-number flows, the added mass matrix can be viewed as an exact representation of the relation between fluid inertial force and body acceleration (32); specifically, *M*_*ij*_ is the added mass coefficient in the *i*-direction corresponding to an acceleration in the *j*-direction; indices 1 to 3 denote linear accelerations along the *x, y*, and *z* axes, whereas indices 4 to 6 represent angular accelerations about these same axes. While this matrix is difficult to determine for bodies of arbitrary shapes, there exist analytical formulae for the added mass coefficients of geometric primitives, drastically simplifying its expression. For this reason, the surfaces of the foot, shank, and thigh were approximated as cylinders following the least squares method described in (33). With its origin placed at the center of the proximal joint, a cylinder aligned with a segment’s long axis *y* admits two planes of symmetry, reducing the number of nonzero, independent added mass coefficients to four. They can then be determined using slender body theory by integrating the 2D coefficients of a circular cylinder of radius *r* over the length of the body *L* (34):

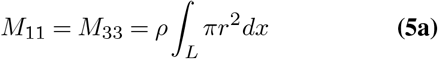

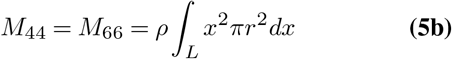

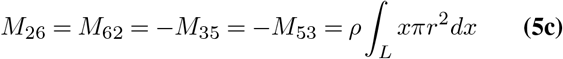

Because axial added mass cannot be determined with the slender body theory, *M*_22_ was approximated using the method of the equivalent ellipsoid (35) and correction factors in (36):

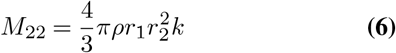

Where

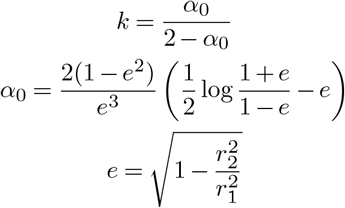

*r*_1_ is the length of the semi-major axis of the equivalent ellipsoid, *r*_2_ is the cylinder’s radius, *e* is the eccentricity of the meridian section, and *α*_0_ reflects the shape of the ellipsoid. Note however that axial added mass is likely of minor importance, as it contributes less than 6% to the total force for slender bodies of fineness ratio > 5 (36)—equivalent to the lower limb geometries in the present study—and axial translation is irrelevant in all exercises but underwater cycling.

Quasi-steady forces acting on the resistive device were calculated likewise. Aquafins were drawn in Blender (37) from images available online, and exported to STL to obtain a triangulated surface onto which local forces can be integrated. As for rectangular plates perpendicular to high-Re flows, lift was ignored. The drag ***F***_*d,aqua*_ is calculated as:

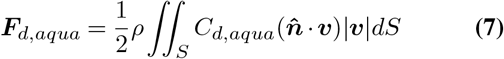

where *C*_*d,aqua*_ is equal to 1.17 (that of a thin plate of aspect ratio ≈2 (38)), and 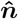 is the local face normal. Added mass was computed with Eqn 4, and the three nonzero coefficients defined in (39) as:

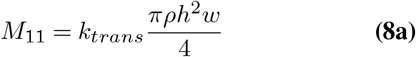

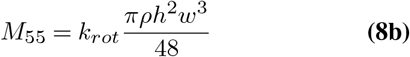

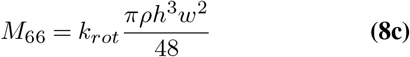

where *k*_*trans*_ and *k*_*rot*_ are correction factors accounting for the finiteness of the Aquafins’ width *w* (0.29 m) and respectively equal to 0.77 and 0.3 (taken from Figs 3 and 4 in (39)), and *h* is the Aquafins’ height (0.14 m).

#### CFD-informed determination of force coefficients

Under the quasi-steady assumption, local forces are uniquely determined by the instantaneous mesh motion; i.e., flow history and transient flow phenomena are implicitly ignored. Therefore, quasi-steady models cannot take into account vortex shedding and wake interaction—two such phenomena that occur during kicking the legs (40)—and also fail to capture the (force-enhancing) proximo-distal axial flow ubiquitous in rotating appendages (41). Furthermore, the hydrodynamics of the human lower limb are poorly documented. With the exception of the experimental study of a foot–shank system by Pöyhönen and colleagues (42), there are no reports of drag and lift coefficients of realistic leg models. To keep the complexity of the model moderate while incorporating otherwise ignored transient hydrodynamical effects, I propose to use high-fidelity numerical fluid flow simulations to derive dimensionless forces at the foot, shank, and thigh. This hybrid approach is computationally inexpensive after calibration, and has proved more accurate than quasi-steady modeling alone in recent flight studies (43, 44).

Fluid flow simulations were carried out with the open-source CFD code OpenFOAM 9 (45). The lower limb geometry was sliced from one (arbitrarily-chosen) full body mesh obtained from Pose2Mesh, cleaned up in Blender, and split into individual foot, shank, thigh and hip patches. The geometry was placed at the center of the computational domain: a rectangular cuboid of length, width, and height 5 × 3 × 5 m, initially coarsely discretized with cells of 0.2 m in all directions using the `blockMesh` utility. This background of hex cells is the prerequisite for achieving finer mesh control with `snappyHexMesh`. The procedure works by iteratively refining the base mesh and morphing it so that it snaps and conforms to the geometry. Specifically, five levels of isotropic refinement were applied over the body surface, as well as inside a cylinder of radius 0.8 m and depth 0.6 m centered on the hip joint^2^. This cylindrical region was added to provide greater resolution of areas of steep velocity and pressure gradients, and, more importantly, to define a region of rotating cells. Mesh motion was prescribed as a square wave function to simulate reciprocating leg motion: movement amplitude was set to 360° with periods *T* of 20, 10, and 6 s (or constant angular velocities of 36, 72, and 120 deg s^−1^); four strokes were simulated. The governing equations for incompressible, turbulent flows with mesh motion were solved using Open-FOAM’s transient solver `pimpleFoam` and the *k*–*ω* SST turbulence model. Foot, shank, and thigh drag and lift coefficients were computed by averaging instantaneous force coefficients from normalized time *t*^*^ = 0.2 to 0.45 and *t*^*^ = 0.7 to 0.95 (i.e., before reversals, when values had stabilized), where *t*^*^ = *t/T* and *t* is the elapsed time. To finely study the relationship between force coefficients and leg orientation, each set of simulations was repeated with the leg at a different angle relative to the incoming flow: geometries were rotated longitudinally from 0° (hip flexion/extension) to 90° (hip abduction/adduction) in steps of 15° (for a total of 21 simulations). Data were then fit using least squares quadratic polynomials to devise predictive formulas for use in our hybrid hydrodynamic model. The discretization error was estimated as recommended in (46) by assessing the convergence of force coefficients computed over three increasingly fine grids (effective refinement ratios of 1.2 and 1.4).

### Prediction of hip and knee joint contact forces

Musculoskeletal simulations were run with OpenSim v4.3 (47) and the high-fidelity, full-body musculoskeletal model created by Rajagopal and colleagues (48). The lower body has 20 degrees of freedom (six describing the pelvis relative to the ground and seven per lower limb), actuated by 80 muscle–tendon units, whereas the upper body has 17 degrees of freedom with a torque actuator each (three at the lumbar joint and seven per upper limb). Muscle model parameters were adjusted to reflect age-related changes in muscle mechanics, with alterations in individual muscle’s isometric strength, maximum shortening speed and normalized force achievable during lengthening, passive tension, and activation dynamics as described in (49). To match the annotated keypoints, 12 virtual markers were added to the model at the origins of the reference frames of the humeri, ulnae, hands, femora, tibiae, and tali. Since this sparse set of markers is insufficient to define all model’s degrees of freedom, 30 additional markers^3^ were defined from the mesh surfaces to prescribe longitudinal rotations and lumbar kinematics more robustly. Body segments were scaled based on average lengths calculated from the 3D poses.

Inverse kinematics (50) was used to solve for the joint angles that place the model in the configuration that best matches the 3D pose from the video; i.e., the posture that minimizes the weighted-least-squares error between the model markers and the corresponding 3D keypoint coordinates at each video frame. Lower weights (*w* = 0.2) were given to the left and right hip keypoints, as these are known to be harder to annotate precisely^4^. Since foot motion was not captured from the videos, the ankle, subtalar, and metatarsophalangeal joints were locked in a neutral position.

Generalized forces (i.e., net joint moments and intersegmental forces) were calculated through inverse dynamics. Kinematic and external force input data were smoothed at the same frequency (second-order, zero-lag, Butterworth filter; low-pass cutoff=3 Hz) to avoid signal artifacts (51, 52), prior to interpolation using generalized cross-validated quintic splines (53) and double differentiation. The muscle redundancy problem (or force-sharing problem; i.e., the indeterminacy caused by the infinitely many ways muscle synergists can contribute to joint moments) was solved using static optimization. The objective was to find the individual muscle forces that produced the joint moments determined by inverse dynamics and minimized the sum of squared muscle activations (a commonly used surrogate for metabolic cost (54)) while constrained by muscle force-length-velocity properties.

Hip and knee joint loads were ultimately evaluated using OpenSim’s `JointReaction` analysis. To match OrthoLoad’s in vivo load recordings, predicted joint forces were normalized to body weight; at the knee, forces were also expressed in subject-specific `PhysicalOffsetFrames` aligned with implant orientation according to specifications at https://orthoload.com/transformation-of-loads. To investigate the relative contribution of individual muscle groups to hip and knee joint loading, joint reaction forces were recomputed one muscle group at a time, prescribing its force to the all-muscle static optimization solution while replacing the remaining muscles with ideal torque actuators, and subtracting the intersegmental force from inverse dynamics (55). For this analysis, only compressive forces (i.e., aligned with the longitudinal axis of the femur or tibia) were considered.

### Statistical analysis

To validate simulations, predicted vs in vivo distributions of force directions were compared using MANOVA owing to its superior statistical power, by linearizing circular data with sines and cosines (56); test statistics were computed using parametric bootstrap based on MATS because of its robustness against covariance matrix singularity and heteroscedasticity (57). Circular statistics were computed in R (58) using the `circular` (v0.4-95; (59)) and `MANOVA.RM` (v0.5.3) packages. One-dimensional Statistical Parametric Mapping—a method using random field theory to make topological inference on smooth continua (originally applied to the analysis of 3D cerebral blood flow, and more recently to biomechanical time series (60))—was done with spm1d v0.4.8 (61). Effects of limb orientation and movement speed on drag and lift coefficients were tested with two-way repeated measures ANOVAs. These and all remaining statistical procedures were performed in `Pingouin` v0.5.2 (62). The significance threshold *α* was set at 0.05.

## Results

Three-dimensional poses and meshes, indispensable inputs to the musculoskeletal models, were predicted from the estimated 2D keypoint coordinates. The average detection error on the 130 unseen images was 1.6 ± 0.2 pixels (equivalent to about 1.1 ± 0.1 cm); errors were largest at the right knee (1.9 ± 1.3 pixels) and smallest at the chin (1.2 ± 0.6 pixels).

To account for unsteady fluid flow effects, our hydrodynamical model was informed by high-fidelity computational fluid dynamics simulations. Following the Grid Convergence Index method (46), the uncertainty due to grid discretization for the dimensionless forces on the lower limb was 3.4%. Hydrodynamic coefficients varied little as limb angular velocity increased from 36 to 120 deg s^−1^ (*F* (2, 4)=0.22, *p*=.70), but were highly dependent on limb orientation (*F* (13, 26)=13.92, *p*=.02; Fig 2). Coefficients at different speeds were therefore pooled and fit by quadratic polynomials so that they could be well approximated with formulas whose coefficients are found in Table 2.

**Table 2.** Coefficients of quadratic polynomials and goodness-of-fit statistics.

| | $a_2$ | $a_1$ | $a_0$ | $r^2$ | MAE |
| --- | --- | --- | --- | --- | --- |
| $C_{d,thigh,0-90}$ | $1.24 \times 10^{-4}$ | $-8.20 \times 10^{-3}$ | $9.02 \times 10^{-1}$ | 0.87 | 0.04 |
| $C_{d,shank,0-90}$ | $7.29 \times 10^{-5}$ | $5.25 \times 10^{-4}$ | $5.65 \times 10^{-1}$ | 0.99 | 0.02 |
| $C_{d,foot,0-90}$ | $-6.85 \times 10^{-6}$ | $4.38 \times 10^{-3}$ | $3.47 \times 10^{-1}$ | 0.88 | 0.03 |
| $C_{l,thigh,0-90}$ | $-3.88 \times 10^{-5}$ | $7.24 \times 10^{-4}$ | $-3.83 \times 10^{-1}$ | 0.83 | 0.03 |
| $C_{l,shank,0-90}$ | $4.04 \times 10^{-6}$ | $-1.02 \times 10^{-3}$ | $-9.26 \times 10^{-2}$ | 0.41 | 0.02 |
| $C_{l,foot,0-90}$ | $2.11 \times 10^{-4}$ | $-2.22 \times 10^{-2}$ | $-7.53 \times 10^{-1}$ | 0.94 | 0.04 |
| $C_{d,thigh,180-270}$ | $-5.27 \times 10^{-5}$ | $2.86 \times 10^{-2}$ | $-2.68$ | 0.85 | 0.05 |
| $C_{d,shank,180-270}$ | $5.23 \times 10^{-6}$ | $2.90 \times 10^{-3}$ | $1.07 \times 10^{-2}$ | 0.95 | 0.03 |
| $C_{d,foot,180-270}$ | $-1.39 \times 10^{-5}$ | $1.27 \times 10^{-2}$ | $-1.61$ | 0.99 | 0.02 |
| $C_{l,thigh,180-270}$ | $-7.49 \times 10^{-6}$ | $-4.63 \times 10^{-4}$ | $1.83 \times 10^{-1}$ | 0.96 | 0.02 |
| $C_{l,shank,180-270}$ | $2.69 \times 10^{-5}$ | $-6.96 \times 10^{-3}$ | $2.92 \times 10^{-2}$ | 0.97 | 0.02 |
| $C_{l,foot,180-270}$ | $1.28 \times 10^{-5}$ | $-1.00 \times 10^{-2}$ | $1.57$ | 0.86 | 0.05 |

Equations of the form $y = a_2x^2 + a_1x + a_0$ were fit to the hydrodynamic coefficients. MAE: mean average error.

**Fig. 2.**
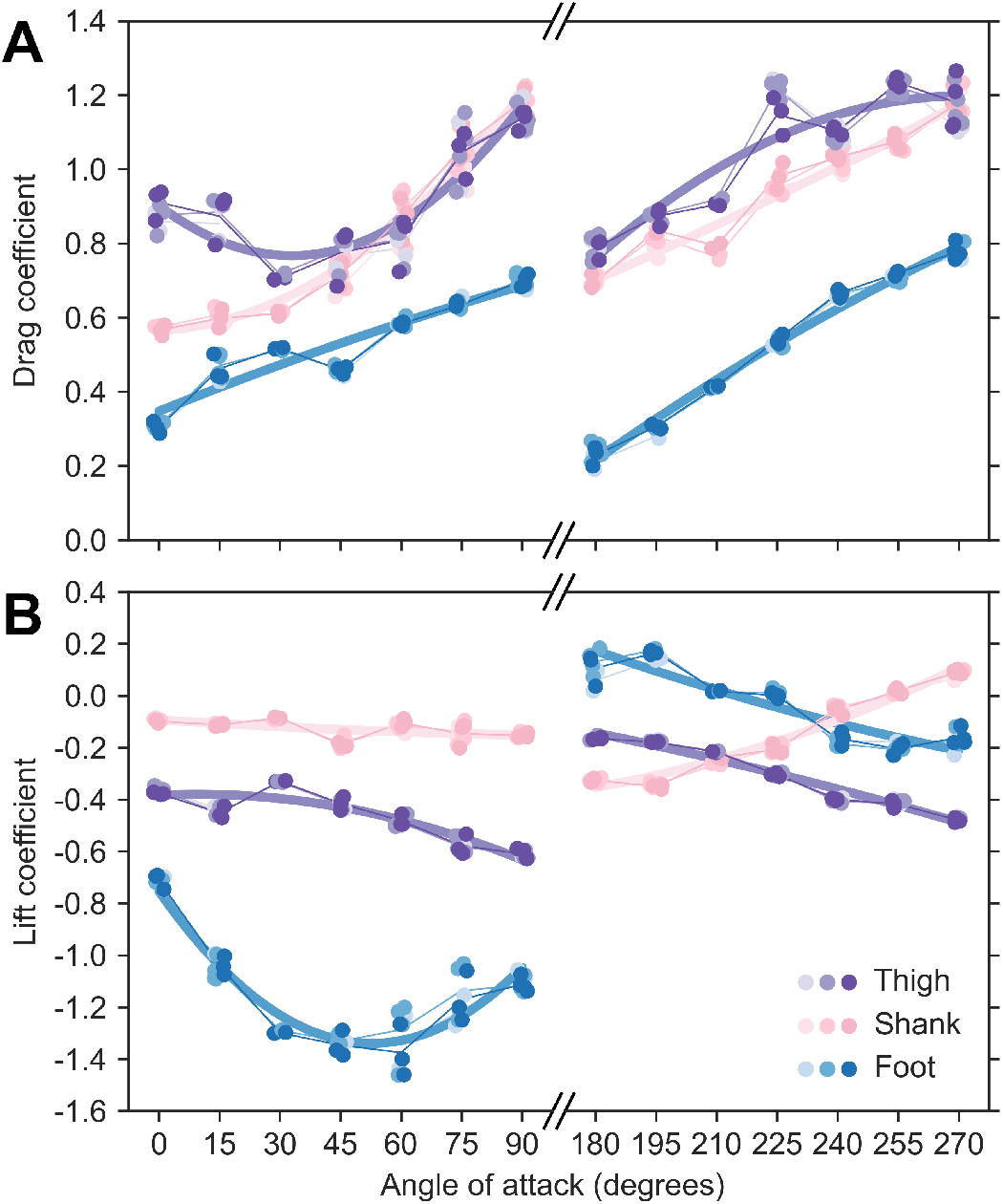
Thigh, shank and foot hydrodynamic coefficients are well-fit by quadratic polynomials. Angle of attack was increased in 15°increments between simulations. Drag (A) and lift (B) coefficients were averaged over the last second of each stroke, when they had stabilized after the transients following start or stroke reversals. Four cycles were simulated at each of the three speeds (36, 72, and 120 deg s^−1^); lighter colors indicate lower speeds. Least squares fits yielded predictive formulas (Table 2) used in the hybrid hydrodynamic model to estimate normal and axial hydrodynamic forces.

Predicted peak joint reaction forces agreed very well with implant recordings. A linear regression analysis revealed that both measurements were strongly correlated (*r*=.80 with 95% confidence interval [.74, .85]; *p*<.001), with a best fit line slope of 0.77 [0.68, 0.86] and intercept of 0.22 [0.10, 0.33] (Fig 3A). Confidence intervals were estimated using bootstrapping and the bias-corrected percentile method. To complement linear regression and further assess the agreement between in vivo and simulated peak joint forces, we used the Bland–Altman analysis (63). Differences between methods were normally distributed (D’Agostino–Pearson test: *K*^2^=3.23, *p*=.20), and no linear trend was apparent, judging from the regression line (slope: –0.11 [–0.35, 0.12]; intercept: 1.13 [1.05, 1.21]; *r*=–.07 [–.23, .10]; *p*=.36). The bias between methods (–0.05 BW) was not statistically significant, as the line of equality was within the confidence interval of the mean difference [–0.10, 0.01]. 95% of the difference fell within limits of agreement extending from –0.71 [–0.80, –0.62] to 0.61 BW [0.51, 0.70] (Fig 3B). When normalized to recorded force peaks, the differences between methods amounted on average to 28.1% BW (range: 15.9–72.0; least in hip flexion, and highest at the knee when cycling).

**Fig. 3.**
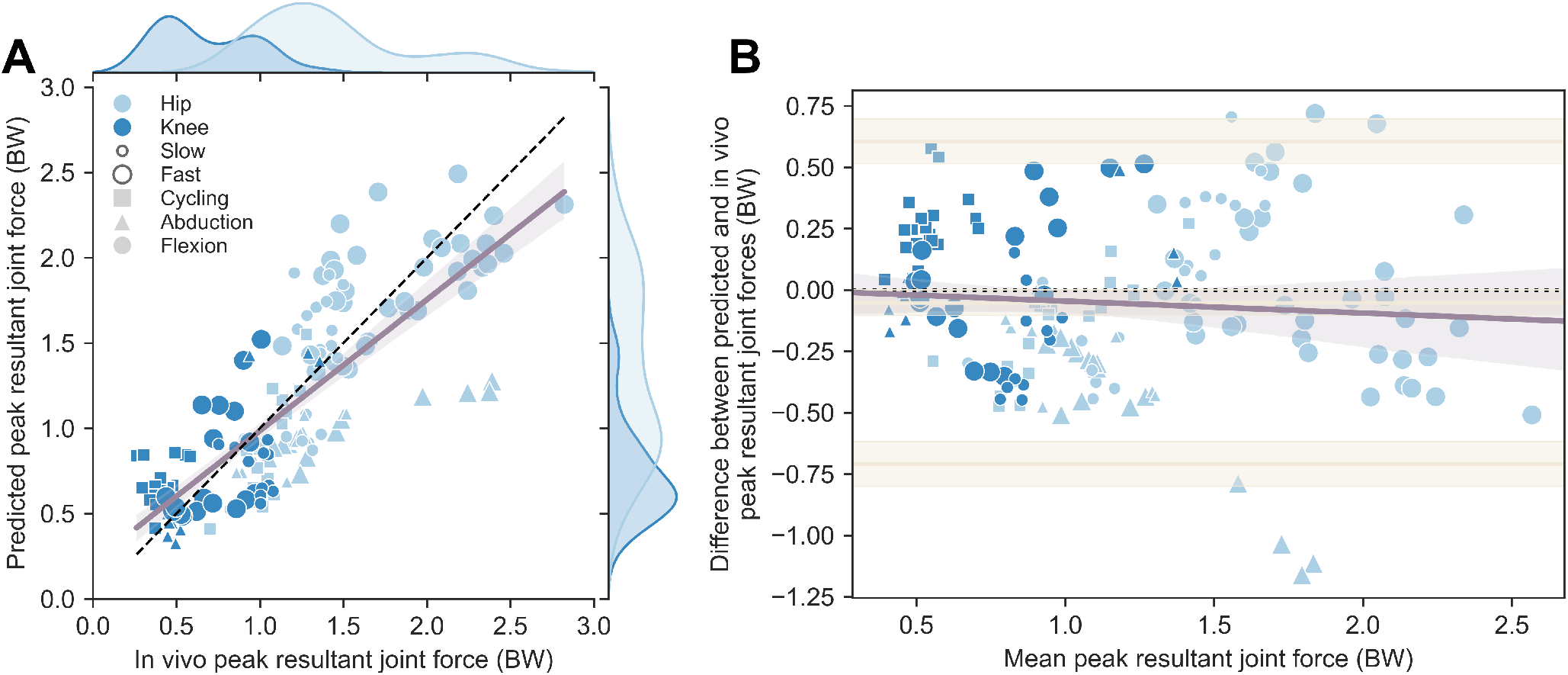
Predicted peak hip and knee joint forces agree well with in vivo implant recordings. Data markers (*N* =160) are colored by joint, with their sizes and symbols changing with exercise speed and type. (A) The black dotted line is the identity line. Also shown is the best linear regression fit and the 95% confidence band estimated using bootstrapping. (B) The Bland–Altman plot reveals good, unbiased agreement between predicted and in vivo peak joint forces, with 95% of the differences found within –0.71 and 0.61 BW.

In addition to force magnitude, we assessed the similarity between predicted and in vivo force directionality. Qualitatively, average directions in the local transverse plane were similar for all exercises and joints but at the knee during cycling (Fig 4A), with angles (relative to the mediolateral axis) and 95% CI of: (i) 136° [135, 137] vs 167° [167, 168]; (ii) –172° [–171, –172] vs 166° [165, 166]; (iii) 141° [140, 141] vs 157° [157, 158]; (iv) –82° [–77, –90] vs 10° [7, 13]; (v) 85° [83, 86] vs 56° [54, 57]; (vi) –71° [–68, –73] vs –73° [–71, –75]. We further tested the null hypothesis that both predicted and in vivo distributions came from the same population (or two populations with the same direction) using a factorial MANOVA design with repeated measures. No significant differences were observed at the hip (cycling: 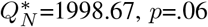; abduction: 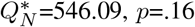 flexion: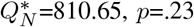) and at the knee during flexion 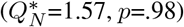, while the null hypothesis was rejected for knee forces during cycling 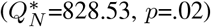 and abduction 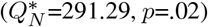. Three-dimensional alignment—as evaluated by cosine similarity (i.e., the dot product between unit vectors; Fig 4B)—was consistently high for both hip and knee joint forces, especially in flexion/extension (hip–slow, as median [interquartile range]: 0.93 [0.91, 0.96]; hip–fast: 0.96 [0.94, 0.97]; knee–slow: 0.94 [0.89, 0.97]; knee–fast: 0.92 [0.89, 0.96]) and abduction/adduction (hip–slow: 0.91 [0.90, 0.99]; hip–fast: 0.95 [0.93, 0.98]; knee–slow: 0.97 [0.96, 0.98]). During cycling, similarity was high at the knee (0.97 [0.92, 0.97]), but lowest at the hip (0.82 [0.65, 0.85]). Collectively, this was equivalent to a median misalignment of 18 [14, 25] degrees.

**Fig. 4.**
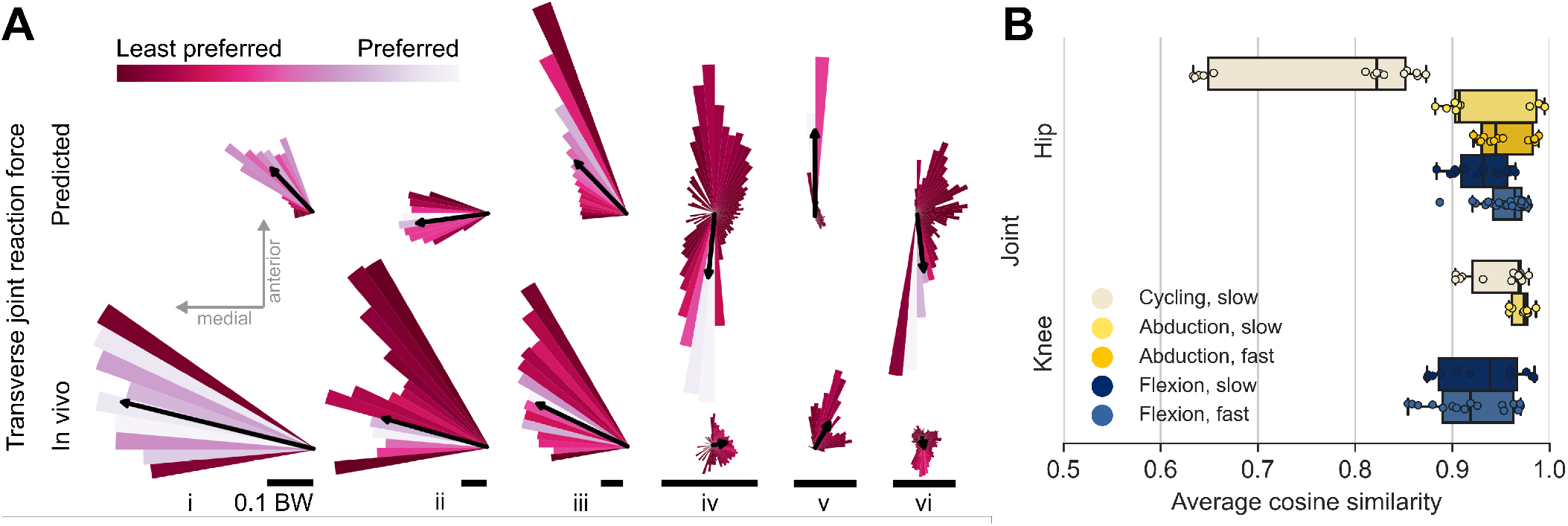
Predicted and measured joint reaction forces are similar in direction in four out of six conditions. (A) Histograms of predicted (top row) and in vivo (bottom) instantaneous force directions in the transverse plane at the hip and knee during cycling (i and iv, respectively), abduction (ii and v), and flexion (iii and vi). Bins (width=5°) of lighter colors reflect the preferred force directions (i.e., most visited angular sectors during the exercise). Black arrows indicate the circular average force direction weighted by magnitude; scale bars represent 10% of the participant’s body weight. Hip joint forces were predicted to be oriented medially and slightly anteriorly, in agreement with implant recordings; simulated transverse forces at the knee swept an angle larger than recorded, though their magnitude remained small (indicating that knee forces are oriented mostly superiorly, in line with the long axis of the tibia). In vivo and predicted angle distributions were statistically similar for all measurements but (iv) and (v). (B) Box plots displaying the cosine similarity distributions between predicted and in vivo 3D force vectors. Whiskers extend to the furthest data point within 1.5× the interquartile range. A similarity of 1 implies perfect vector alignment, whereas a value of 0 indicates orthogonal vectors. With the exception of hip force vectors during cycling, simulated forces were aligned within ≈ 15° with in vivo forces.

Instantaneous muscle activations (Fig 5) were determined using static optimization—a procedure aiming at resolving joint moments from inverse dynamics into individual muscle forces while satisfying muscle contraction dynamics. Significant statistical differences in activation between speeds were determined using statistical parametric mapping and are shown as darker regions in the 95% CI bands. The vastus lateralis was the least activated muscle across all exercise modalities. With the exception of a short burst of activity about 65% of the cycle when using Aquafins, the hamstrings and gluteus maximus remained silent during slow hip extension. Fast hip flexion/extension was achieved not only through a significantly greater activation of the prime movers (notably when exercising with the resistive device), but also by a marked inhibition of the iliopsoas, rectus femoris, and sartorius during hip extension. A similar behavior was observed during hip adduction, where higher speed was met by the greater recruitment of the adductors together with a decreased activation of the gluteus medius and the gastrocnemius. No significant differences were seen during knee flexion/extension. Cycling exhibited moderate activation of the hip flexors, but high activation of the gastrocnemius, hip extensors, and gluteus medius, with peak activity exceeding that during hip extension with Aquafins for the latter two.

**Fig. 5.**
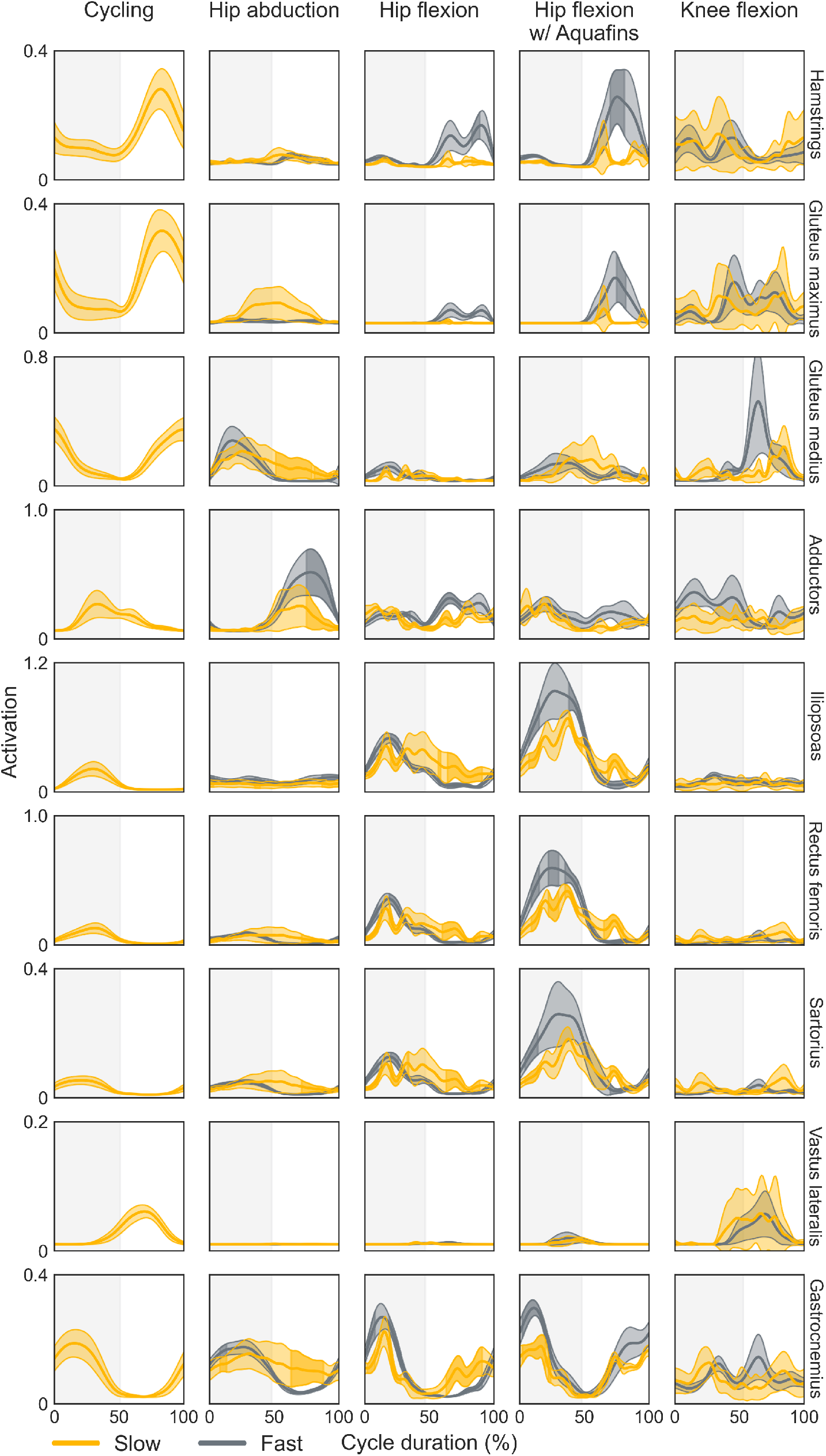
Higher exercising speeds at the hip are met by increased activity of the prime movers and inhibition of the antagonists. Average instantaneous lower limb muscle activations (thick lines) and 95% confidence bands during slow (yellow) and fast (gray) exercises are shown. Darker band regions indicate significant differences in activation between speeds, as determined via statistical parametric mapping. Shaded gray vertical areas denote cycle reversals (e.g., the transition from hip abduction to adduction, or from knee flexion to extension); note that cycling begins with the recovery phase. Higher exercising speeds at the hip were accompanied by increases in activation of the prime movers (e.g., hip flexors/extensors during hip flexion/extension, most notably when wearing the Aquafins), whereas no significant differences were observed during knee flexion/extension. Some muscles were grouped according to their functions, and their activations summed: hamstrings (biceps femoris long and short heads, semimembranosus, semitendinosus); adductors (adductor brevis, longus, and magnus); iliopsoas (psoas major and iliacus); gastrocnemius (medial and lateral heads). Although the musculoskeletal model comprises more muscles, only the nine most relevant are displayed here.

Prescribing a subset of muscles to their forces from static optimization while substituting the remaining ones with ideal torque actuators allowed us to elucidate the contribution of individual muscle groups to joint contact forces. Hip flexors, glutes, adductors, and hamstrings were the main contributors to hip joint compressive forces during hip-driven exercises (40.4±12.7%, 25.6±9.7%, 14.2±4.8%, 13.0± 8.2%, respectively). The gastrocnemius and vasti only contributed 5.1±2.6% and 1.9±0.5% (Fig 6); conversely, they produced 39.1±15.9% and 29.4±13.7% of the knee compressive forces, followed by the hamstrings (15.4±11.3%), the hip flexors (14.1± 8.2%), glutes (1.8±1.6%), and adductors (0.3±0.6%). Changes in the contribution of individual muscle groups to a joint’s compressive forces accompanying increases in movement speed could be examined further for knee flexion, as well as hip abduction and flexion. Only during fast hip flexion did the compressive force increase significantly (0.73±0.21 BW, *p*<.001; Fig 7A). This was mainly caused by the larger forces produced by the hip flexors (0.56±0.15 BW, *p*<.001), the gluteal muscles (0.12±0.07 BW, *p*<.001), and the gastrocnemius (0.05±0.01 BW, *p*<.001). Compressive forces otherwise decreased at high speed during hip adduction (–0.20±0.15 BW, *p*=.003) as a result of lower glutes (–0.14±0.10 BW, *p*=.04) and hip flexors (–0.08±0.06BW, *p*<.001) forces, and during knee extension (–0.28±0.23 BW, *p*<.001) due to reduced gastrocnemius (–0.13±0.11 BW, *p*<.001) and hip flexors (–0.09±0.06 BW, *p*<.001) forces (Fig 7B).

**Fig. 6.**
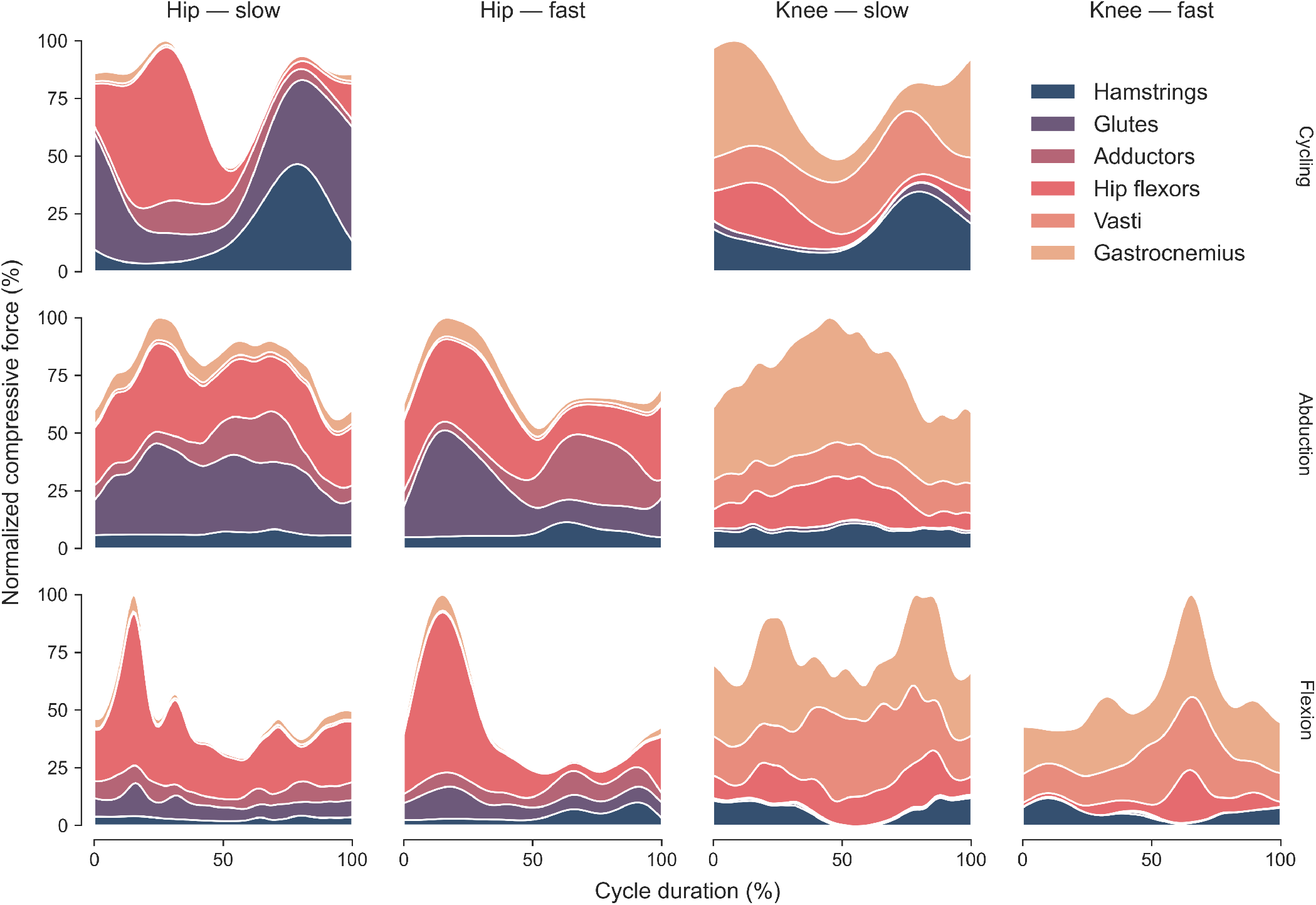
Hip flexors and gastrocnemius contribute the most to hip and knee joint compressive forces. The hip flexors were the main contributors to hip compressive forces, particularly during hip flexion and the recovery phase of cycling, followed by the glutes and the adductors. The hamstrings contributed only little compression during all exercises but the power phase of cycling. The contribution of the gluteal muscles and adductors to knee compressive forces was close to null, whereas the gastrocnemius and the vasti muscles were dominant. Similar to Fig 5, muscles were grouped based on their functional roles: glutes (gluteus minimus, medius, and maximus); hip flexors (iliacus, psoas major, rectus femoris, and sartorius); vasti (vastus medialis, intermedius, lateralis).

**Fig. 7.**
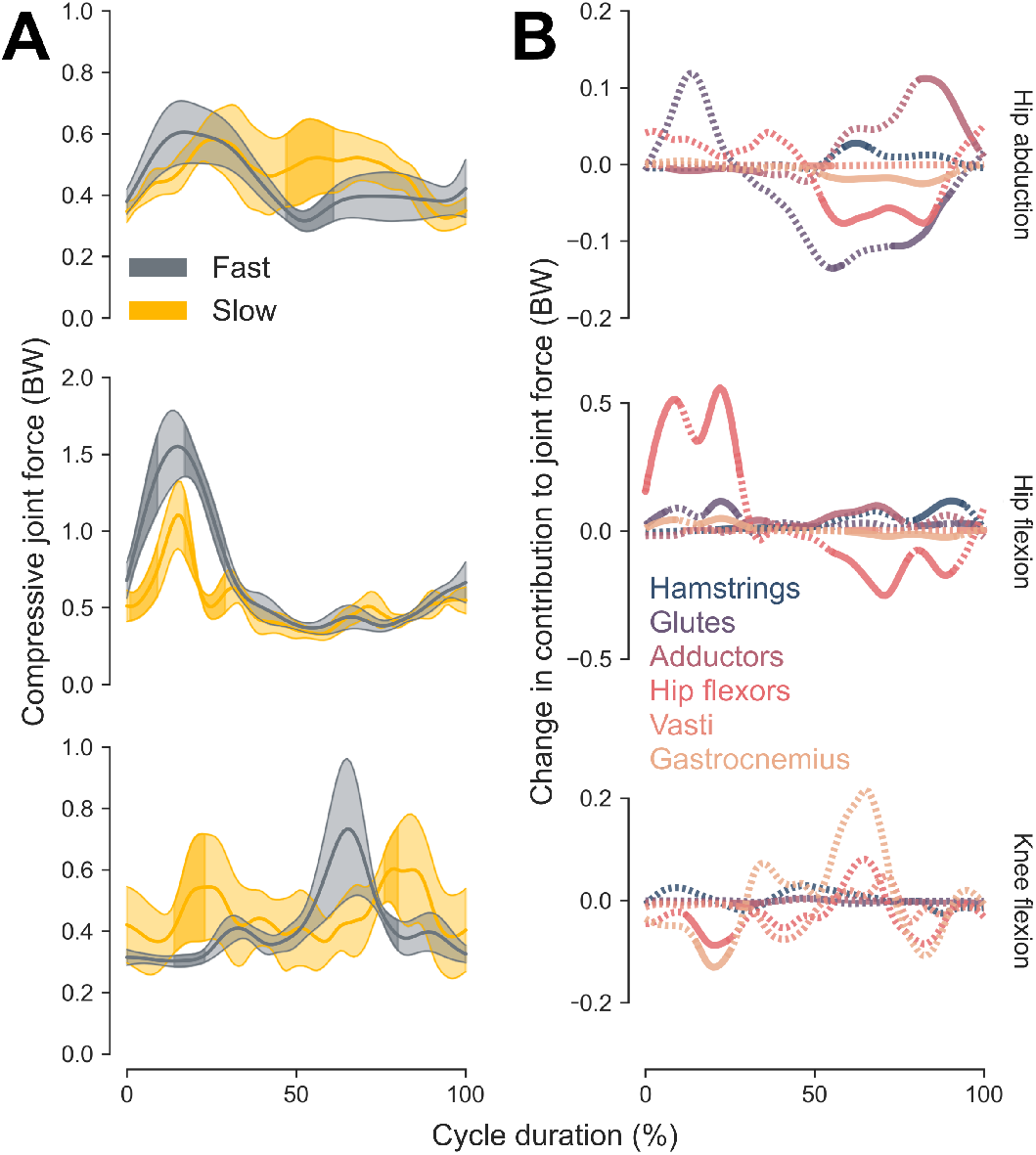
Changes in contribution to joint compressive forces as speed increases are attributed primarily to the hip flexors and gluteal muscles. (A) Hip forces were noticeably larger during fast hip flexion (0.6 BW; darker regions in the 95% CI bands; middle row), while they significantly decreased during knee extension (bottom) and towards movement reversal during hip abduction (top). (B) Corresponding changes in joint force as speed increased, decomposed into individual muscle group contribution. Solid lines represent statistically significant changes. Decreased hip forces during hip adduction were caused mainly by a reduction in glutes (–0.14 BW) and hip flexors (–0.08 BW) forces, whereas decreased knee forces were the result of lower gastrocnemius (–0.13 BW) and hip flexors (–0.09 BW) forces. Larger hip joint compression during fast hip flexion was attributable to the hip flexors (0.56 BW), the glutes (0.12 BW), and, to a lower though significant extent, the gastrocnemius (0.05 BW).

## Discussion

I described a markerless method to predict lower limb joint loading during aquatic exercises from a single video only. This noninvasive and inexpensive approach—bringing together new developments in computer vision, computational biomechanics, and hydrodynamic modeling—automatically estimates an individual’s three-dimensional pose and body surface, the forces exerted on the water, and ultimately the forces produced by muscles and those of contact between joints. The latter was strongly correlated with in vivo recordings from instrumented implants (*r*=.80). 95% of the differences in magnitude between methods were found within –0.71 and 0.61 BW, an error range comparable with the most accurate predictions of previous gait modeling studies (64). The present solution thus offered an accuracy equivalent to simulations driven by laboratory-grade motion capture systems, but with a single-camera setup.

Joint force directions too were well predicted. The cosine similarity between simulated and in vivo 3D force vectors was consistently high, and the distributions of forces along the anteroposterior and mediolateral axes were statistically indistinguishable from implant recordings in four out of six conditions. Shear forces are specifically crucial to understanding joint stability or pain etiology (e.g., at the hip (65)). In the absence of tissue modeling, they are also commonly used proxies for ligament or cartilage strains (e.g., for the anterior cruciate ligament (66)). As such, accurately determining force direction in silico will likely prove key to the informed selection of safe aquatic exercises, and the improvement of rehabilitation outcomes eventually.

Non-hip- and non-knee-spanning muscles had little effect on joint loading, contributing on average less than 7.0% and 2.1% to hip and knee compressive forces, respectively. This is unlike dry-land locomotion (e.g. (67, 68)), during which some muscles have considerable influence on the loading of joints they do not anatomically cross. This phenomenon, known as dynamic coupling (69), is dependent on a system’s inertia and arises due to the redirection of external forces by muscles and the transmission of forces between segments. In water, it is perhaps attenuated by buoyancy and added mass—which makes segments virtually heavier—though further analysis is warranted to clarify muscle contributions to hydrodynamic forces. Two important clinical consequences follow from the above: across the exercises simulated here, joint compressive forces are most effectively modulated by the prime movers (particularly, the hip flexors and gastrocnemius); other muscles are not protective, as they have minimal to no offloading effect.

Thanks to our novel CFD-informed hydrodynamic model, the time to compute hydrodynamic forces was cut down from hours (using high-fidelity numerical fluid flow simulations) to seconds while preserving relevant transient effects. Furthermore, preliminary experiments trading shape recovery accuracy for speed by approximating body parts as a set of articulated geometric primitives (e.g., truncated cones for the thighs and shanks) hint at an additional order of magnitude reduction in processing time. Jointly with the use of DeepLabCut-Live! SDK (70), this opens the way for realtime processing of video streams, with potential for the delivery of biofeedback; e.g., to reduce joint loading in postoperative patients, or to reach some water resistance threshold for conditioning in athletes.

Some limitations ought to be acknowledged. First, our biomechanical simulations relied on scaled, albeit generic musculoskeletal models. Low geometric specificity is known to entail adverse effects on joint kinematics and load estimates (71–73). Automated and fast subject-specific reconstruction of lower limb skeletal models is nowadays possible, but still necessitates manual processing of body scans (74). A notable yet budding alternative resides in the direct estimation of the skeleton from an individual’s body shape, circumventing the need to acquire medical images; however, further effort is warranted to make it comply with biomechanical standards (75). Second, static optimization inherently underestimates coactivation. Because the objective was to minimize the sum of squared muscle activations, it is muscles with long moment arms and large torque-generating capacity that are preferentially recruited, at the expense of muscles with smaller leverage (76). Incorporating muscle synergies into the optimization formulation may yield more realistic antagonistic activation (77), although it requires prior knowledge about the synergy structure. Provided that experimental electromyographic data are available, the cost function could also be adjusted (78). Nonetheless, small and deep hip muscles were found to have insufficient ability to produce stiffness (79). Therefore, inadequately estimating its function was unlikely to have critically impacted joint loading prediction. Third, Pose2Mesh imperfectly recovers body shape to the extent that poses contain limited cues about corpulence or leanness (29). While visually modest in the present study, error is expected to increase in obese or athletic populations, contaminating the calculation of hydrodynamic forces and in turn the prediction of joint- and muscle-level kinetic quantities. To mitigate this issue, additional shape cues could come from part segmentation supervision (e.g., using differentiable semantic alignment loss (80)), or networks specifically designed to regress shape from images and leverage sparse body measurements (81).

## Conclusion

I proposed an accurate and noninvasive method to predict lower limb joint loading in water from monocular videos, combining recent advances in markerless mesh recovery, hydrodynamic modeling, and gold-standard musculoskeletal simulations. The computation of mechanical demands enables the standardization and clearer prescription of an exercise’s intensity, and is expected to improve dissemination and reproducibility of rehabilitation protocols. Finally, it lays the ground for predictive simulations (82); e.g., for the design of aquatic equipment or the discovery of new movements satisfying individualized task goals.

## Acknowledgments

I wish to warmly thank Asim Iqbal for early stimulating discussions, as well as João Paulo Vilas-Boas, Johan Lambeck, Maxime Vidal, and Aristotelis Economides for critical feedback.

## Footnotes

1 An accessible overview of the topic is presented in (24).

2 Given that refinement is defined relative to the background mesh size and that every level splits a cell in eight new ones, this resulted in 2^3*5^ = 32, 768 new cells of edge length 0.2*/*2^5^ ≈ 0.006 m for every original hex in these regions.

3 calcaneus; heads of the first and fifth metatarsals; medial and lateral malleoli, femoral and humeral epicondyles; ulnar and radial styloids; anterior and posterior superior iliac spines; incisura jugularis; processus xiphoideus; C7; and, T10.

4 see §2.2 at https://cocodataset.org/#keypoints-eval.

